# Automated design of highly diverse riboswitches

**DOI:** 10.1101/603001

**Authors:** Michelle J Wu, Johan O L Andreasson, Wipapat Kladwang, William Greenleaf, Rhiju Das

**Affiliations:** Program in Biomedical Informatics, Stanford University, Stanford, CA, USA; Department of Genetics, Stanford University, Stanford, CA, USA; Department of Biochemistry, Stanford University, Stanford, CA, USA; Department of Applied Physics, Stanford University, Stanford, CA, USA; Department of Physics, Stanford University, Stanford, CA, USA

**Keywords:** riboswitch, RNA, molecular design, high-throughput measurements, thermodynamic model, computer-assisted design

## Abstract

Riboswitches that couple binding of ligands to recruitment of molecular machines offer sensors and control elements for RNA synthetic biology and medical biotechnology. Current approaches to riboswitch design enable significant changes in output activity in the presence vs. absence of input ligands. However, design of these riboswitches has so far required expert intuition and explicit specification of complete target secondary structures, both of which limit the structure-toggling mechanisms that have been explored. We present a fully automated method called RiboLogic for these design tasks and high-throughput experimental tests of 2,875 molecules using RNA-MaP (RNA on a massively parallel array) technology. RiboLogic designs explore an unprecedented diversity of structure-toggling mechanisms validated through experimental tests. These synthetic molecules consistently modulate their affinity to the MS2 bacteriophage coat protein upon binding of flavin mononucleotide, tryptophan, theophylline, and microRNA miR-208a, achieving activation ratios of up to 20 and significantly better performance than control designs. The data enable dissection of features of structure-toggling mechanisms that correlate with higher performance. The diversity of RiboLogic designs and their quantitative experimental characterization provides a rich resource for further improvement of riboswitch models and design methods.

## Main text

Biological systems rely on precise regulation of cellular processes. In particular, regulatory RNAs, including riboswitches, play major roles in biological circuits, sensing molecules in the cellular milieu and then modulating gene expression and other processes in a wide variety of natural systems.^1^ The ability to perform *de novo* design of arbitrary riboswitches that interact with other biomolecules in their environments would have broad impacts in synthetic biology as well as for RNA diagnostics and therapeutics. Supporting these efforts, there are a rapidly growing number of synthetic and natural RNA ‘aptamer’ sequences that bind drugs, metabolites, proteins, and other biologically important molecules that might be incorporated into novel riboswitches. Many applications of these riboswitches, including fluorescent biosensors,^2–6^ require reversible riboswitches with tight binding to reporters in their ON states, and this criterion necessitates a tradeoff with good activation ratios, defined as the ratio in observed signal in the presence and absence of a trigger molecule.^7^

Riboswitches are multi-stable RNA molecules, meaning they can form multiple secondary structures. The preferred states can be toggled by small molecule inputs or RNA oligonucleotides that bind aptamers or complementary regions embedded in the RNA (Figure 1A). So far, the majority of riboswitch design studies involve manual design of the desired states and require detailed specification of the structure-toggling mechanism.^7^ For reversible switches, these efforts have required significant trial-and-error; success has been achieved through screening of many constructs, the majority of which exhibit little to no switching, with median activation ratios close to 1 and best-case activation ratios of 10.^2,3,6,7^

**Figure 1:**
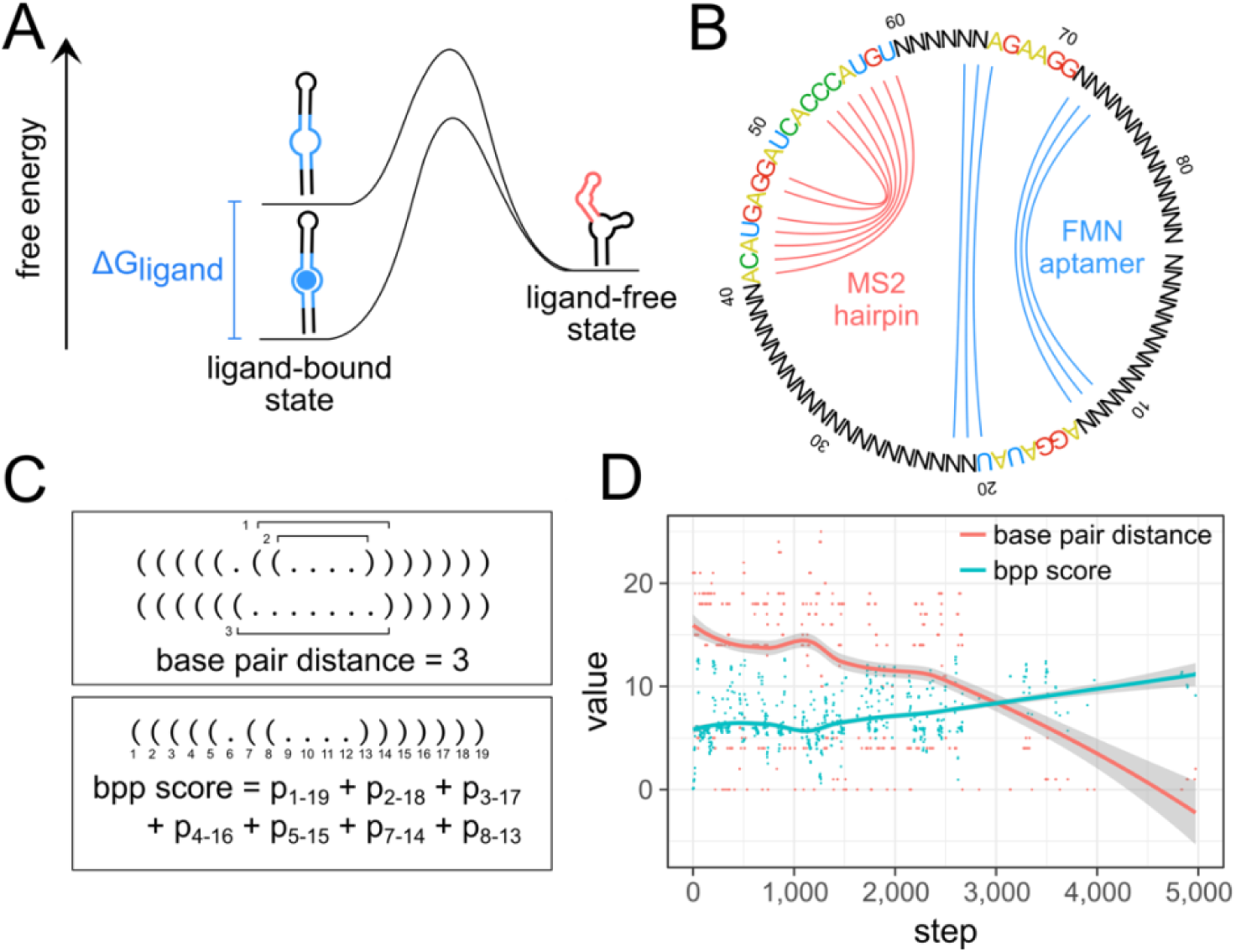
RiboLogic uses a graph representation and two scoring functions to design riboswitches. (A) This energy diagram represents the thermodynamic model used, where the ligand-bound state is given an energetic bonus due to the chemical potential of the binding of the ligand. (B) A graph representation is used to constrain the sequence space that is sampled by RiboLogic. In this example, the goal is to design a riboswitch whose formation of the MS2 RNA hairpin is modulated by the presence of the flavin mononucleotide (FMN) molecule. Bases connected by an arc are part of these secondary structure elements and are constrained to be complementary in sequence update. (C) Two scoring metrics are used to evaluate each design candidate. The base pair distance measures the number of base pairs that must be broken or formed to reach the target structure, while the base pair probability (bpp) score quantifies the probability of formation of each base pair in the target structure. (D) The scores change as expected during computational design, with the base pair distance decreasing and the base pair probability score increasing over optimization steps.

Proposals to automate this process still require experts to input the complete secondary structures of the RNA without and with ligands or are limited to specialized applications.^8–13^ In addition to lack of generality, lack of diversity, and limited automation, most methods have been subjected to no experimental tests^8–11^ or tests involving at most tens of riboswitches.^12,13^ It remains unclear if successful riboswitches can be created without expert input of detailed secondary structures and to what extent current energetic models for RNA secondary structure or expert-defined structure-toggling mechanisms might limit more automated efforts.

Here, we present a detailed computational and experimental study involving thousands of diverse molecules to test the fully automated design of riboswitches. For computational design, we describe RiboLogic, an algorithm for designing sequences of RNA molecules that are predicted to change their secondary structure in response to interactions with other biomolecules. This package only requires the user to provide small aptamer segments to bind desired input and output molecules. For experimental characterization, we evaluate the switching of thousands of designed RNA molecules using repurposed Illumina sequencers, through the recently developed the RNA-MaP (RNA on a massively parallel array) platform.^14–17^ These experimental results confirm that fully automated design can yield riboswitches with performance comparable to rational design, achieving activation ratios above 10 in many cases. The large number of measurements and high diversity of structure-toggling mechanisms allow dissection of currently limiting factors for automated riboswitch design and provide a rich data set for future efforts that seek to improve riboswitch design through machine learning or more accurate physics-based modeling.

RiboLogic designs riboswitches based on a maximally flexible set of user-specified constraints. The algorithm accounts for any number of folding conditions, as defined by the concentrations of ligands defined by the user. These ligands can be small molecules, proteins with known aptamers, or other RNA strands engaged through base-pairing interactions. For example, in some of our tests below, we used flavin mononucleotide (FMN) as an input ligand; FMN binds to a small aptamer sequence discovered by *in vitro* selection (Figure 1A & 1B).^18^ The user only needs to specify the sequence of this aptamer and the estimated dissociation constant of the aptamer-ligand complex under the experimental conditions, and RiboLogic will place this ‘input’ segment within the design and optimize the surrounding sequence in each of the riboswitch states, simulating ligand binding to the aptamer (see Methods for details). In this example, the two states are RNA with no FMN present and with a concentration of 200 μM FMN (Figure 1A). For each of the target riboswitch states, the user can specify either a full desired secondary structure or, more simply, the substructure of an ‘output’ segment that must be adopted or not adopted by the RNA in order to trigger or suppress an output, respectively. For example, in some of our tests below, we used binding of a fluorescently tagged MS2 viral coat protein to an MS2 RNA hairpin segment within the design as an output (Figure 1A & 1B); such interactions underlie most systems for CRISPR interference and activation and *in situ* RNA visualization.^2–6,19–21^ The user only needs to specify the sequence and ‘active’ secondary structure of this output element and RiboLogic will place this sequence relative to the input aptamer element and optimize surrounding sequences during its design process.

RiboLogic uses simulated annealing to sample the space of possible sequences to satisfy the given constraints. At each step, the sequence is mutated either at a single base or by sliding the position of a functional element (e.g., the FMN aptamer or MS2 hairpin; colored nucleotides in Fig. 1B). For each sequence that is sampled, the minimum free energy secondary structure is determined for each solution condition (e.g., without and with 200 μM FMN) and evaluated by two scores (Figure 1C & 1D). The first score is a base pair distance that measures the number of base pairs that must be broken or formed to obtain the target structure or substructures in each solution condition, summed over the different solution conditions. The second score is a base pair probability score that sums the probabilities of formation of all base pairs that should form in the target structure or substructures, providing a smoother quantitative measure of structure formation than the first base pair distance score. RiboLogic implements several additional strategies to narrow the sequence space being explored. Mutation of the sampled sequences leverages a dependency graph-based approach, which ensures that bases that are paired in any target structure are always complementary in sequence (e.g., N’s connected by blue lines in Figure 1B).^22^ In the case of designing riboswitches responsive to other input RNA molecules, the algorithm provides the option to automatically introduce the sequence complementary to the input in order to promote favorable interactions between the designed RNA and input RNA.

As test cases for our methods, we designed riboswitches where the binding of a small molecule or oligonucleotide ligand modulates the formation of the MS2 RNA hairpin, which can then transduce outputs by recruiting machinery coupled to the MS2 bacteriophage coat protein.^23–25^ We applied a quantitative, high-throughput array technology that enables fluorescence measurements over millions of individual RNA clusters generated on an Illumina array, which has been extensively tested using the MS2 system (Figure 2A & 2B).^14,16,17^ The formation of the MS2 RNA hairpin was detected by flowing fluorescently labelled MS2 protein at increasing concentrations to get a binding curve (Figure 2B & 2C). The dissociation constant *K*_*d*_ was fit over tens to hundreds of clusters for each design, yielding a distribution of *K*_*d*_ measurements for each state (Figure 2D). By taking the median of each distribution, we calculated a *K*_*d*_ as a quantitative measure of the switching of each design, and the ratio of these MS2 *K*_*d*_ values with and without input ligand (e.g., FMN) gives an activation ratio, which we use as our figure of merit for riboswitches. This activation ratio is equal to the ratio of fluorescence of the riboswitch with and without input ligand at low MS2 concentrations^7^; by carrying out fits of data from sub-nanomolar to many micromolar MS2 concentrations, we achieve high precision in these measurements. The resulting *K*_*d*_ values and activation ratios were strongly correlated across experimental replicates, confirming the high precision of the method (*r*^2^=0.94 for log *K*_*d*_; errors in activation ratios well under 2-fold; see Figure S1).

**Figure 2:**
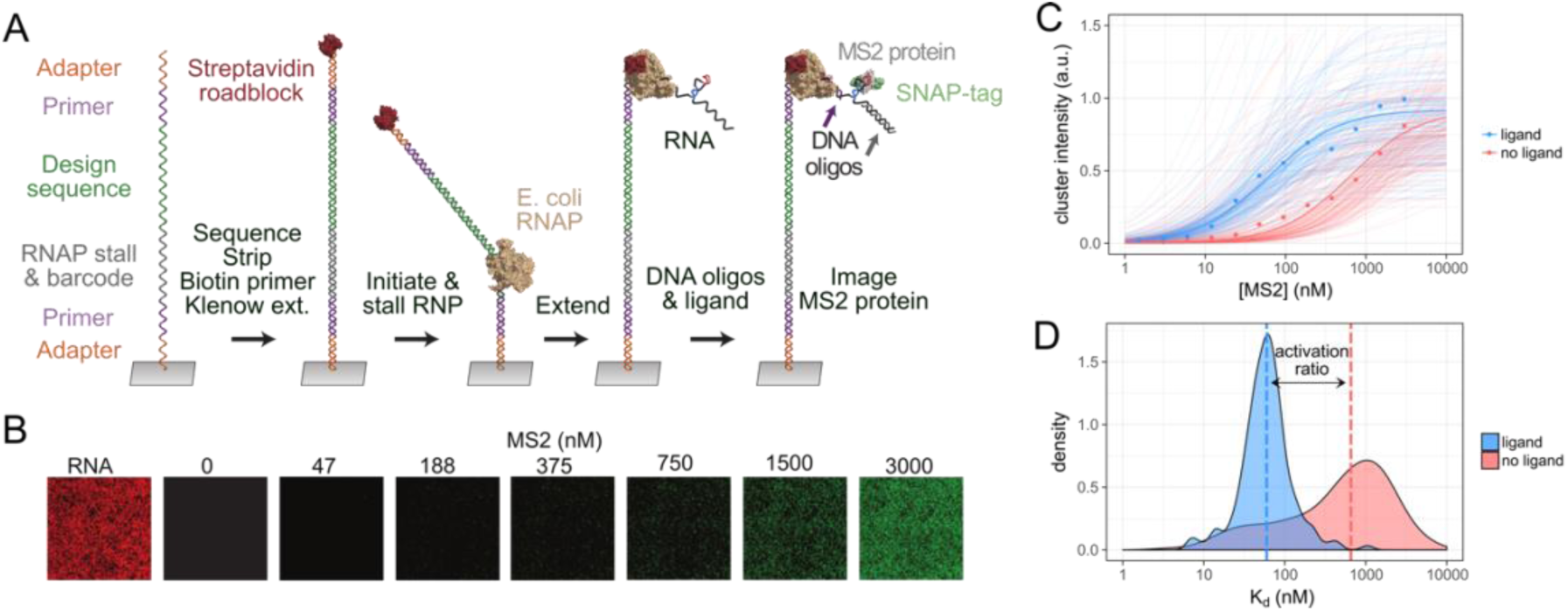
Functional tests of riboswitches using a high-throughput array. (A) Each cluster on the array initially contained a single species of ssDNA from a synthesized oligo pool. dsDNA was generated by Klenow extension with a biotinylated primer, and RNA was transcribed by RNA polymerase until being stalled at the streptavidin roadblock. (B) Fluorescently-labelled MS2 protein was flowed in at varying concentrations to enable measurement of binding. (C) The array technology enables measurement of binding curves over tens or hundreds of replicate clusters for each design and solution condition. (D) The median over the distribution of fit *K*_*d*_s was used to estimate the activation ratio of switching. In this example of an ON switch, the activation ratio of 11 was measured over 172 independent clusters displaying the same switch.

We applied the algorithm to design simple switches responsive to three different small molecules – flavin mononucleotide (FMN), theophylline, and tryptophan. For OFF switches, the MS2 hairpin should form when the ligand is absent and be disrupted when the ligand is added (Figure 3A). For ON switches, the MS2 hairpin should form only when the FMN is present and otherwise be disrupted (Figure 3B). By applying secondary structure constraints to the MS2 hairpin region in both the absence and presence of the ligand, we set up a simple two-state design problem. We were able to obtain a set of structurally diverse designs (Figure 3A-C), and we experimentally characterized thousands of these molecules with the RNA-MaP method.

**Figure 3:**
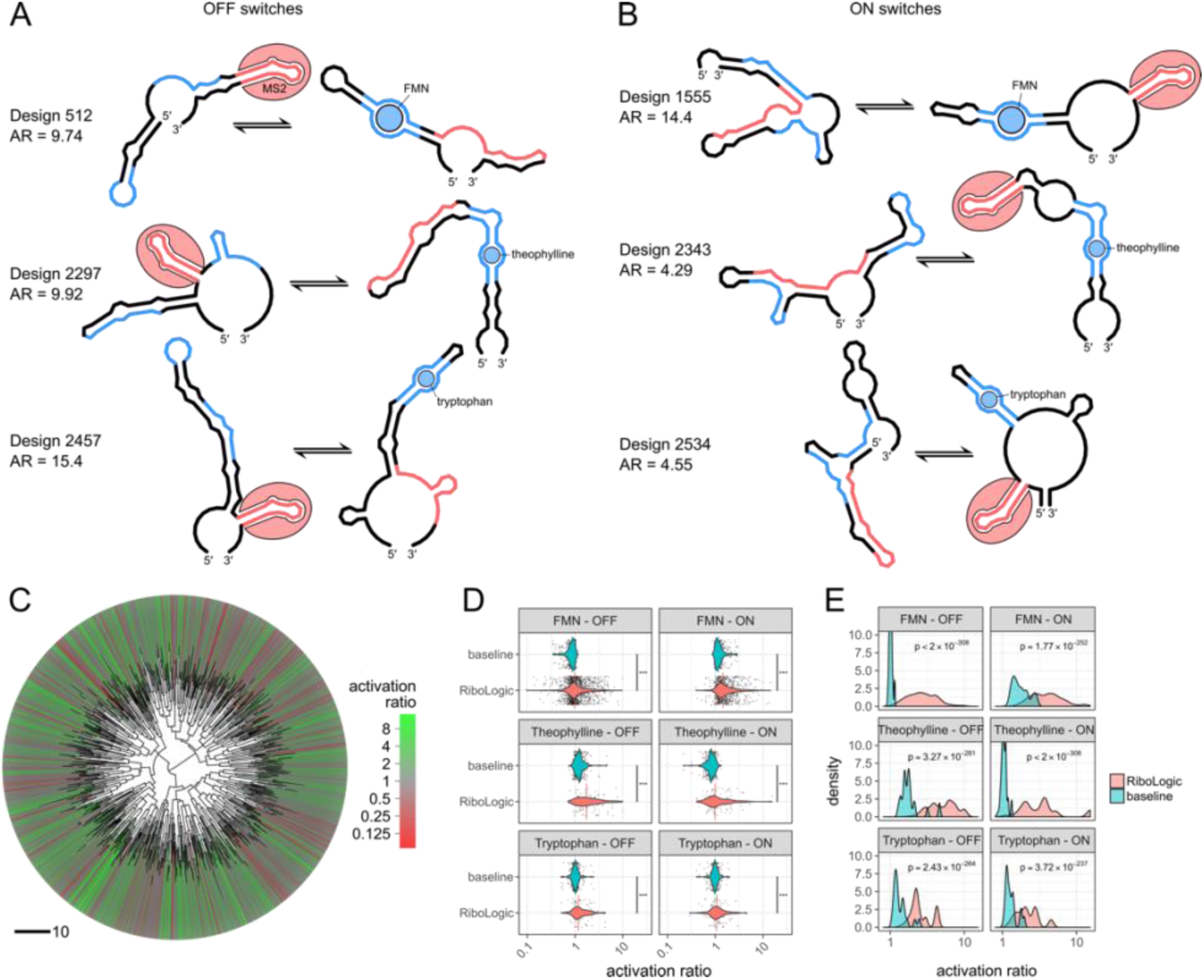
Design of ligand-responsive riboswitches. (A) Predicted secondary structures for a variety of OFF switches show disruption of the MS2 hairpin (red) upon binding of FMN, theophylline, or tryptophan (blue). (B) Predicted secondary structures for a variety of ON switches show formation of the MS2 hairpin (red) upon binding of FMN, theophylline, or tryptophan (blue). (C) Clustering of FMN switches based on the sum of base pair distances of predicted secondary structures reveals that RiboLogic designs with diverse structures achieve high activation ratios. (D) Distributions of experimentally measured activation ratios are shown for various types of designs, with medians shown as vertical lines. RiboLogic generally achieves significantly better activation ratios than baseline, as determined by a Wilcoxon rank-sum test (*** −p<0.001). Baseline is the measured activation ratio for sequences made for other design problems. (E) In practice, several of the most promising designs would be experimentally screened to evaluate switch efficiency. To mimic this, we bootstrapped sets of ten designs and chose the design with the best activation ratio. The distributions of activation ratios for these best-of-ten designs were compared between RiboLogic and baseline. A best-of-ten strategy yields designs with significantly higher activation ratios than baseline.

We found that RiboLogic designs achieved activation ratios significantly better than unrelated designs made for other ligands, which were used as baseline comparisons (Figure 3D). For example, the median activation ratio for Ribologic designs of FMN-responsive ON switches was 1.5 with a standard deviation of 1.3 (Figure 3D, Table 1, Table S1). As the baseline comparison, the median activation ratios with respect to FMN for designs meant to be responsive to theophylline or tryptophan was 1.2. For each of the six switch design challenges (three ligands, ON vs. OFF) the difference was significant (p<10^−10^; Figure 3D, Table S2). For comparison, previous characterization of rationally designed reversible riboswitches yielded a median activation ratio of 1.1.^3,6^

**Table 1:** Summary of activation ratios for RiboLogic designs.

| design | maximum AR | median AR | best-of-ten median AR | count |
| --- | --- | --- | --- | --- |
| FMN OFF | 9.74 | 0.987 | 2.57 | 1357 |
| FMN ON | 14.4 | 1.46 | 3.89 | 853 |
| theophylline OFF | 9.92 | 1.73 | 4.86 | 97 |
| theophylline ON | 15.4 | 0.991 | 3.44 | 99 |
| tryptophan OFF | 4.29 | 1.17 | 2.28 | 89 |
| tryptophan ON | 4.55 | 1.08 | 2.09 | 94 |
| miRNA OFF | 21.8 | 0.825 | 1.66 | 188 |
| miRNA ON | 20.0 | 1.17 | 2.84 | 98 |

For each of the six challenges, the best activation ratio was over 4-fold, and extended up to 15-fold for the theophylline ON switch tests (Figure 3D). Anticipating that most riboswitch design efforts will be able to experimentally test several molecules and choose the best one, we conducted a best-of-ten analysis, in which we randomly drew subsets of 10 designs and scored the best activation ratios. These best-of-ten trials showed clear separation of the activation ratios from baselines, and in the majority of cases gave activation ratios of 2.0 or greater (Figure 3E, Table S3). In addition, most designs exhibited *K*_*d*_’s close to the affinity of the MS2 coat protein under the conditions in which they were supposed to be active (with ligand for ON switches; without ligand for OFF switches) (Figure S2). The switch with the highest activation ratio of 15.4 achieved a *K*_*d*_ of 10 nM in the activated state, within experimental error of the intrinsic dissociation constant of the MS2 coat protein-RNA hairpin interaction (6 nM, measured in the same experiment).

We further tested if RiboLogic could design riboswitches that are responsive to RNA inputs instead of small molecule ligands. Specifically, we applied the algorithm to design 286 switches that modulate MS2 binding based on the presence of miR-208a, a 22-nt miRNA implicated in cardiac hypertrophy.^26^ This type of RNA-based system could be used in diagnostic devices or linked to downstream therapeutic events. Using RiboLogic, we were able to design both ON and OFF switches triggered by the miRNA strand (Figure 4A & 4B). We found that these designs generally took more iterations of optimization to satisfy the constraints as compared to the ligand-responsive switches (Figure S2), but diverse mechanisms were achieved (Figure 4C). While experimental evaluation showed no significant difference between RiboLogic and baseline designs in terms of the median activation ratio, the best-of-ten comparison showed significant differences and maximum activation ratios of 20 exceeded those of small molecule activated switches (Figure 4D & 4E, Table 1). These computational and experimental observations suggest that design for RNA-responsive switches may be intrinsically more difficult, despite the larger binding energy of the RNA compared to the small molecule ligands, perhaps due to a large number of competing binding modes where the input RNAs hybridize to alternative locations in the riboswitch design. At the same time, this automated procedure can still lead to excellent microRNA sensors at the expense of characterizing more designs.

**Figure 4:**
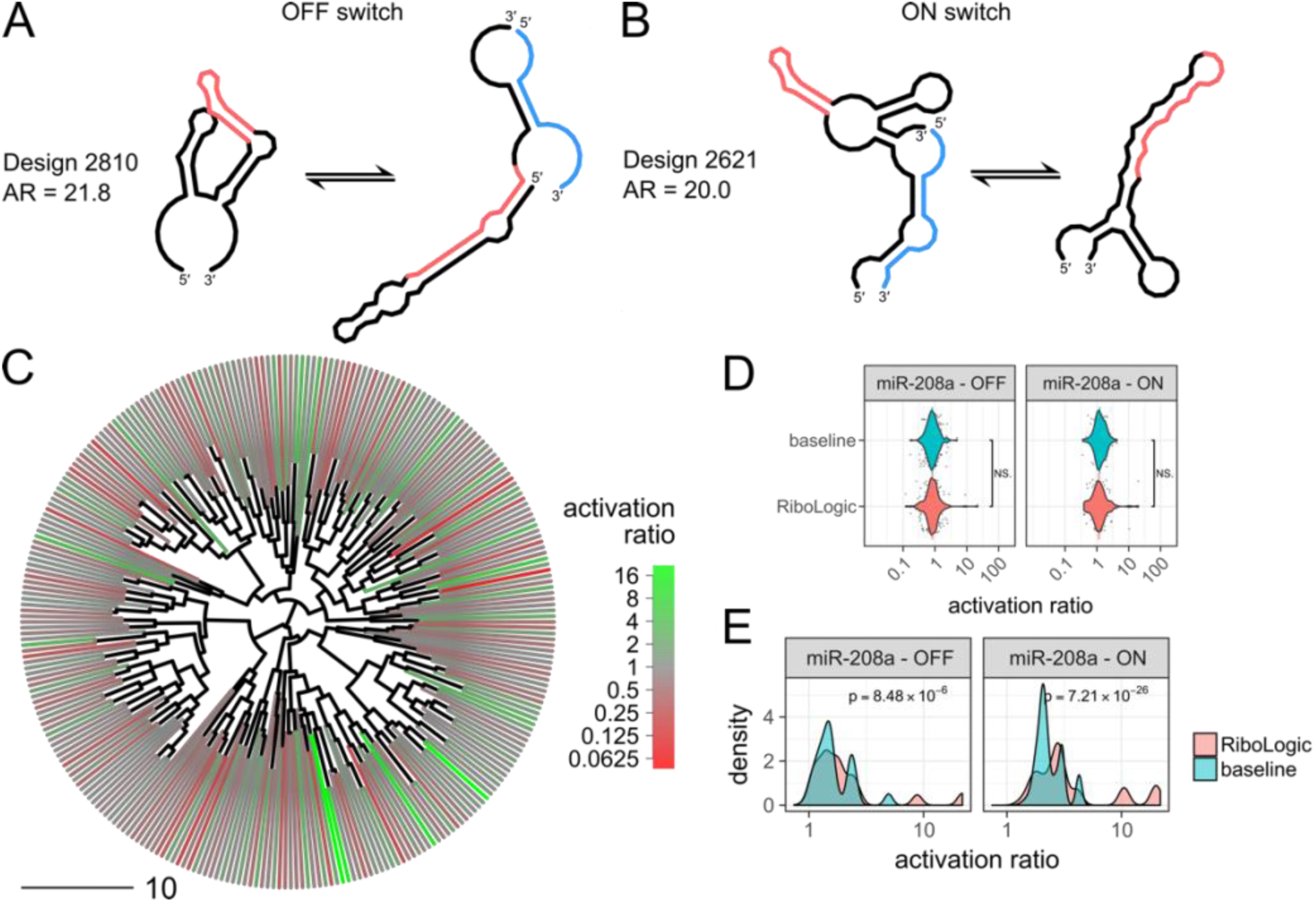
Design of miRNA-responsive riboswitches. (A) This OFF switch is predicted to form the MS2 hairpin (red) only in the absence of the miRNA (blue). (B) This ON switch is predicted to form the MS2 hairpin (red) only in the presence of the miRNA (blue). (C) Clustering of miRNA switches based on the base pair distance between predicted secondary structures in the absence of the miRNA reveals that RiboLogic designs with diverse structures achieve high activation ratios. (D) The distribution of experimentally measured activation ratios are shown as scatter and violin plots, with medians shown as horizontal lines. Across all design problems, there is no significant difference between RiboLogic and baseline designs, as determined by a Wilcoxon rank-sum test. (E) A best-of-ten strategy analysis results in designs with significantly higher activation ratios, but the distributions are similar with the exception of a few outliers.

Across these design challenges, we found that riboswitches with high activation ratios could take a variety of forms. Some high performing designs had the MS2 sequence nested between the two sides of the aptamer, while others had the MS2 outside, with only a short hairpin between the two halves of the ligand-binding internal loop (Figure 3A & 3B; compare designs 2297 & 1555 to 512 & 2534). Some designs formed relatively simple secondary structures with long stems, while others formed more complex folds with three-way junctions (Figure 3A & 3B; compare designs 512 & 2357 to 1555 & 2534). Several structures contain large single-stranded regions, while some have regions designed to bind the functional elements when they are inactive (Figure 3A & 3B; compare design 2534 to 512). The size of our dataset enabled detailed analyses of these secondary structure features, highlighting several that were significantly correlated with high activation ratios (Figure S4). For example, the data showed that having more base pairs shared between states correlated with higher resulting activation ratios. Still, the correlations of any single feature with activation ratio are weak (*r*^2^ < 0.5). Regression models that take into account multiple features will be interesting to develop and test.

A related insight into current design limitations is also enabled by the diversity and large number of our riboswitches. We note that the designs produced by RiboLogic have features that are distinct from designs created by human experts. For the small molecule sensitive riboswitches (Figure 3), the RiboLogic designs include numerous stems outside the aptamer segments that need to be broken or formed. These designs are not as ‘concise’ as expert-designed riboswitches seen in the literature^2,12^, although it should be noted that some natural riboswitches do involve ornate conformational rearrangements.^27^ For the miRNA-sensitive riboswitches (Figures 4), the binding of the input miRNA and the RiboLogic riboswitch is typically not through a completely contiguous, long RNA-RNA duplex, as is typically the case in, e.g., toehold riboswitches^28,29^ or DNA logical devices^30,31^ designed by human experts. Automated riboswitch design might improve if these features seen in human designs were rewarded or seeded into the RiboLogic design algorithm.

We hypothesized that errors in current RNA secondary structure energetic models might be limiting for RiboLogic designs. We carried out comparisons of *K*_*d*_’s and activation ratios predicted by the ViennaRNA and NUPACK packages for small molecule and miRNA riboswitches, respectively. We saw poor correlations for both (*r*^2^ of 0.06 and 0.01 for small molecule and miRNA riboswitches, respectively; Figure S5 & 5). Several designs predicted to have poor activation ratios (near or lower than 1.0) in fact gave activation ratios near 10.0; and other designs predicted to have outstanding activation ratios (greater than 100.0) gave experimental activation ratios lower than 1.0 (Figure 5B). This experiment-theory correlation was better for small-molecule riboswitches compared to the miRNA riboswitches, consistent with the generally better activation ratios of the former, relative to baseline measurements (compare Figures 3 and 4; Table S1). Future design efforts would benefit from more accurate computational models of RNA folding energetics; we present all data collected herein in Supplemental Data to help guide and validate such improvements.

**Figure 5:**
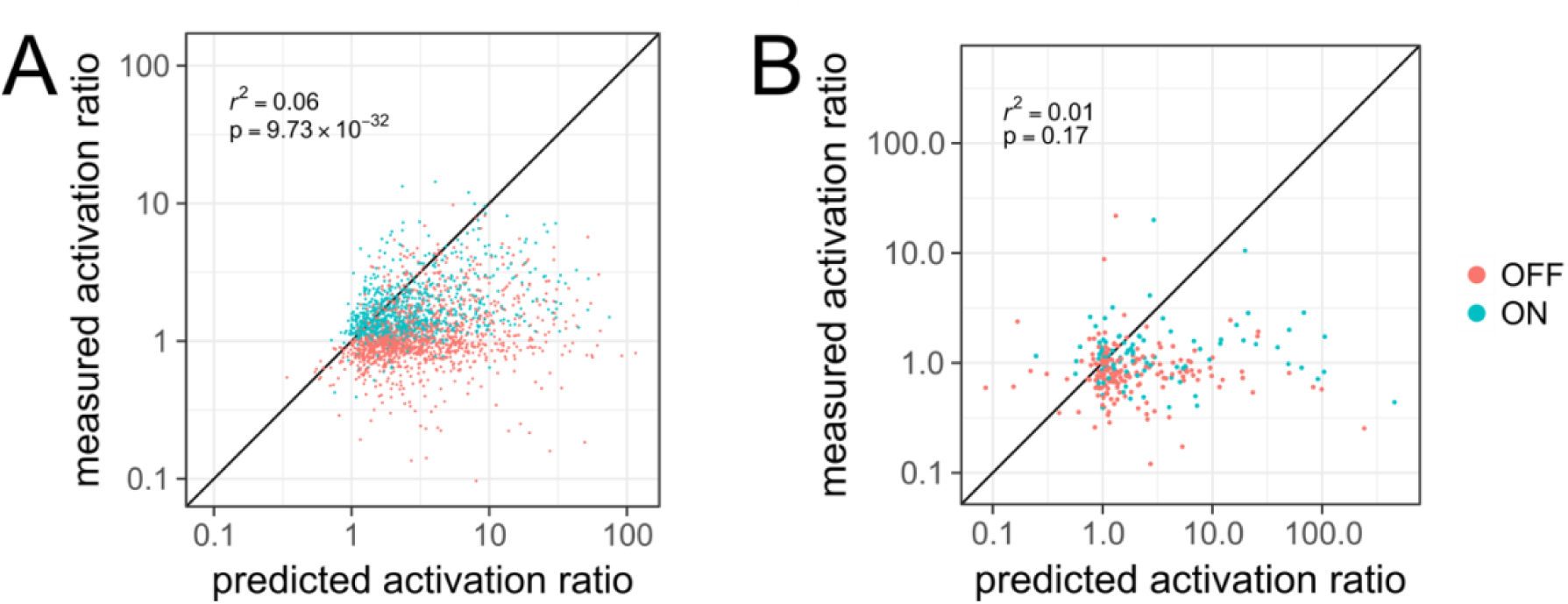
Comparison of predicted and measured activation ratios. (A) For small molecule riboswitches, the predicted activation ratio is somewhat correlated with measured activation ratio. (B) For miRNA riboswitches, the correlation between prediction and experiment is poor.

Here, we have presented RiboLogic, an automated algorithm for designing riboswitches, as well as a rich dataset characterizing a few thousand ligand-responsive RNAs. We show that RiboLogic generates designs with diverse structural mechanisms and achieves activation ratios comparable to previous efforts in rational design of reversible riboswitches. In combination with improved thermodynamic models and high-throughput measurement techniques, we expect that this method and these data will enable improved automated design of switchable RNA elements for a wide variety of applications in biotechnology and medicine.

## Methods

### Design algorithm

#### Overview

Given secondary structure constraints in multiple states defined by ligands or short RNA inputs, our method optimizes an RNA sequence using a simulated annealing algorithm. The starting sequence is selected to ensure complementarity in the target secondary structures. In each step, a random mutation is made, and the new sequence is evaluated using a base pair distance and a base pair probability score. The sequence is updated based on a Metropolis-Hastings acceptance criterion:

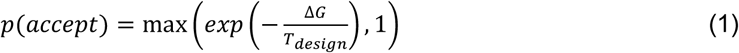

where Δ*G* is the difference in score between the updated and current sequences and *T*_*design*_ is the temperature parameter. This temperature parameter is decreased over the course of the optimization and can be tuned by the user. By default, it decreases linearly from 5 to 1 over the course of design. This process is repeated until a satisfactory sequence is found or the maximum number of iterations specified by the user is reached.

#### Constraints

Sequence constraints can include fixed bases at specified positions as well as substrings that are disallowed from the final sequence. Secondary structure constraints can be given for multiple user-specified states, as defined by varying concentrations of the input ligands. For small molecule and protein ligands, the aptamer sequence, secondary structure, and dissociation constant must be specified. For each state, secondary structure constraints can be applied to any part of the input sequence, including any RNA inputs, and bases can be specified to be unpaired, paired to any other base, or paired with a specific other base. Secondary structure elements’ positions can be left unspecified, and RiboLogic will optimize its position as well. To further ensure diversity, for the tests herein, we enforced two different global arrangements of the aptamer and MS2 hairpin elements – one with the two parts of the aptamer loop adjacent to each other and one with the MS2 sequence nested within the aptamer segments.

#### Sequence update

Sequences are represented in a dependency graph structure as described by Flamm *et al.*^22^ Briefly, each base is a node and each base pair in the constraints forms an edge between nodes. The graph is maintained such that nodes connected by an edge are always complementary. Each time a base is mutated, its entire connected component is mutated accordingly to ensure that all nodes connected to the selected base maintains complementarity. In addition, sequence constraints are incorporated into this graph, disallowing mutations that would force a constrained base to change. In the case of RNA inputs, our method provides the option to automatically introduce the complement of the input sequence into the design sequence in order to promote interactions between strands. This complementary segment can be altered in length, moved, or mutated as a sequence update step.

#### Scoring functions

Two scoring functions are used: a primary score based on a single minimum free energy secondary structure, and a base pair probability-based secondary score that is used in the primary score’s place when the it is the same between two sequences. Based on the predicted minimum free energy structures in each state, a base pair distance to the target secondary structure is calculated. The base pair distance is the number of base pairs that must be broken or formed in order to get from one secondary structure to the other.^32^ If only a substructure is specified, this can include the breaking of base pairs formed with nucleotides outside of the subsequence specified. In addition, for small molecule riboswitches, if the energy of the ligand-bound conformation, with energetic bonus, is not lower than the ligand-free conformation, a penalty equal to the Δ*G* between the two states is applied to the base pair distance.

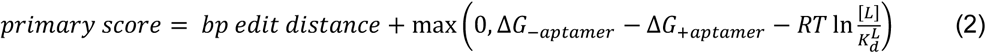

where Δ*G*_*-aptamer*_ is the free energy of the RNA alone in kcal/mol, [*L*] is the concentration of the input ligand, 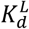 is the affinity of the input ligand, Δ*G*_*+aptamer*_ is the free energy of the RNA constrained to form the aptamer, is the gas constant, is the experimental temperature (37 °C = 310.15 K). We consider only structures that form the desired aptamer, as opposed to doing a minimum free energy calculation with an energetic bonus. This allows the algorithm to guide the sequence towards those that have a more favorable aptamer-forming conformation, even if it is not the minimum free energy structure. We used a value of 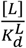 of 133 for FMN and 150 for theophylline and tryptophan, based on initial *K*_*d*_ estimates for those input ligands and experimental [*L*] = 200 μM, 2 mM, and 2.4 mM (FMN, theophylline, and tryptophan, respectively).

However, since the score in eq. 2 is not highly sensitive to single mutations, a secondary base pair probability score is used when the base pair distance is unchanged between sequence updates. This measure of secondary structure formation over the full ensemble is defined by

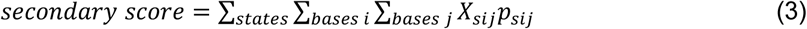

where *s* is the index of the folding state, *i* and *j* are indices of the base position in the sequence, *X_sij_* is an indicator variable representing whether base *i* and *j* should be paired in state *s*, and *p*_*sij*_ is the probability of base *i* and *j* forming in state *s* according to the partition function calculation. The value of the indicator variable is 1 if the base pair should be formed, −1 if it should not be formed, and 0 if it is unconstrained.

Folding of each sequence can be modeled using either ViennaRNA^33^ or NUPACK.^34^ NUPACK 3.0.5.^34^ was used for design involving more than one RNA, in order to properly model multi-strand RNA folding, while ViennaRNA 2.1.9^33^ was used for designs involving small molecule aptamers.

The score used for the Metropolis-Hastings criterion in eq. 1 was:

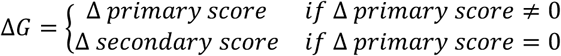

### Computation and code availability

All computation was performed on Intel Xeon Processors E5-2650. The code is available at https://github.com/wuami/RiboLogic.

Average computation time for the design of a ligand-induced riboswitch varied widely, both across runs and depending on the design problem (Figure S3). Every 1,000 iterations took about 2 minutes on one core.

### High-throughput array experiments

The experimental methods have been described in detail previously^14,16^. Briefly, DNA templates for designs were synthesized (CustomArray, Bothell, WA) and sequenced on Illumina MiSeq instruments, and RNA was transcribed directly on the sequencing chip in a repurposed Illumina Genome Analyzer II instrument. Fluorescently-labelled MS2 protein was introduced at concentrations from 1.5 nM to 3 μM, and fluorescence images were collected and quantified to generate binding curves in buffer of 100 mM Tris-HCl, 80 mM KCl, 4 mM MgCl_2_, 0.1 mg/ml BSA, 1 mM DTT, 10 μg/ml yeast tRNA, 0.012% Tween20. These curves were measured in the absence and presence of the ligand of interest, with concentrations of 200 μM FMN, 2 mM theophylline, 4 mM tryptophan, and 100 nM miR-208a. These conditions were selected based on the *K*_*d*_ of each ligand. Each design was measured over an average of about 100 individual clusters on the flow cell. Median fit *K*_*d*_ values over all clusters for each design were used to compute the activation ratio. Designs were prepared and analyzed as part of the Eterna massive open laboratory experiments (rounds R95, R101, and R107).

Designs for which *K*_*d*_ measurements were made over fewer than 10 clusters were excluded from our analysis to avoid poor quality measurements. For diversity analysis, Levenshtein distance was computed between each pair of sequences to obtain a distance matrix. Complete-linkage hierarchical clustering was performed to obtain a dendrogram with each design as a leaf (hclust in R). For statistical analysis, two-sided Wilcoxon rank sum test was used to determine if activation ratios between design types were significantly different. Predicted *K*_*d*_’s were computed as described by Wayment-Steele et al.^7^ Calculations were performed in R^35^, with example scripts available at https://github.com/wuami/RiboLogic. The full dataset is available as Supplementary Data.

## Acknowledgments

We thank F. Portela, J. Anderson-Lee, E.Fisker, and R. Wellington-Oguri for discussions of these designs. This work was funded through a Burroughs-Wellcome Foundation Career Award (to RD), NIH Grant R01 GM100953 (to RD), Stanford School of Medicine Discovery Innovation Award (to RD), and a JIMB Seed Grant (to RD and WJG). MJW was supported by NSF Graduate Research Fellowship DGE-114747, NLM Biomedical Informatics Training Grant T15 LM007033, and NIH Ruth L. Kirschstein National Research Service Award F31GM125151. Computational design was performed on the Stanford BioX3 cluster, supported by NIH Shared Instrumentation Grant S10 RR02664701.

