## Supplementary Information for "Automated design of highly diverse riboswitches"

### Figure S1

**Reproducibility of experimental measurements.** $K_d$ (A) and activation ratio (B) values measured over two replicates correlate with an $r^2$ (in log space) of 0.94 and 0.84, respectively. The color represents the minimum number of clusters across the two replicates. The dotted lines denote the boundary for error within a factor of 2 between the two measurements.

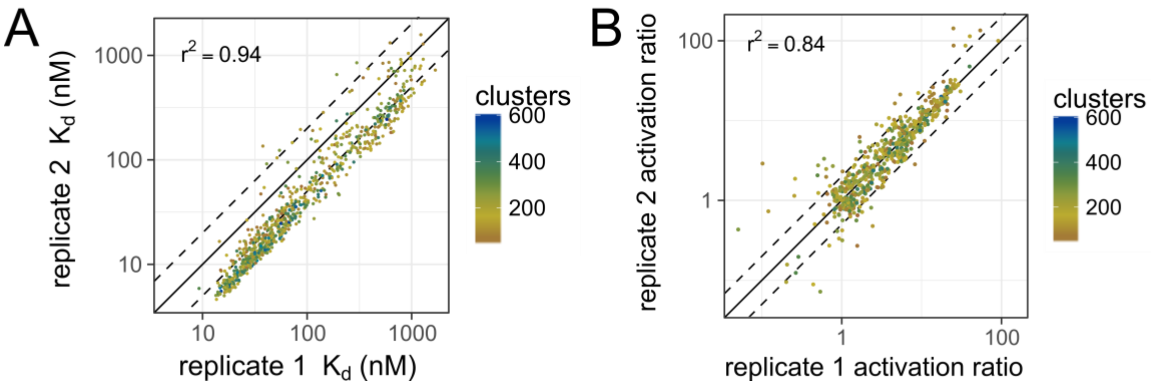

### Figure S2

**On and off state $K_d$'s for RiboLogic small molecule riboswitches.** $K_d^{ON}$ vs. $K_d^{OFF}$ plots show that most designs achieve $K_d^{ON}$ within a factor of 10 of the intrinsic $K_d$ under conditions where they should be activated (dotted lines).

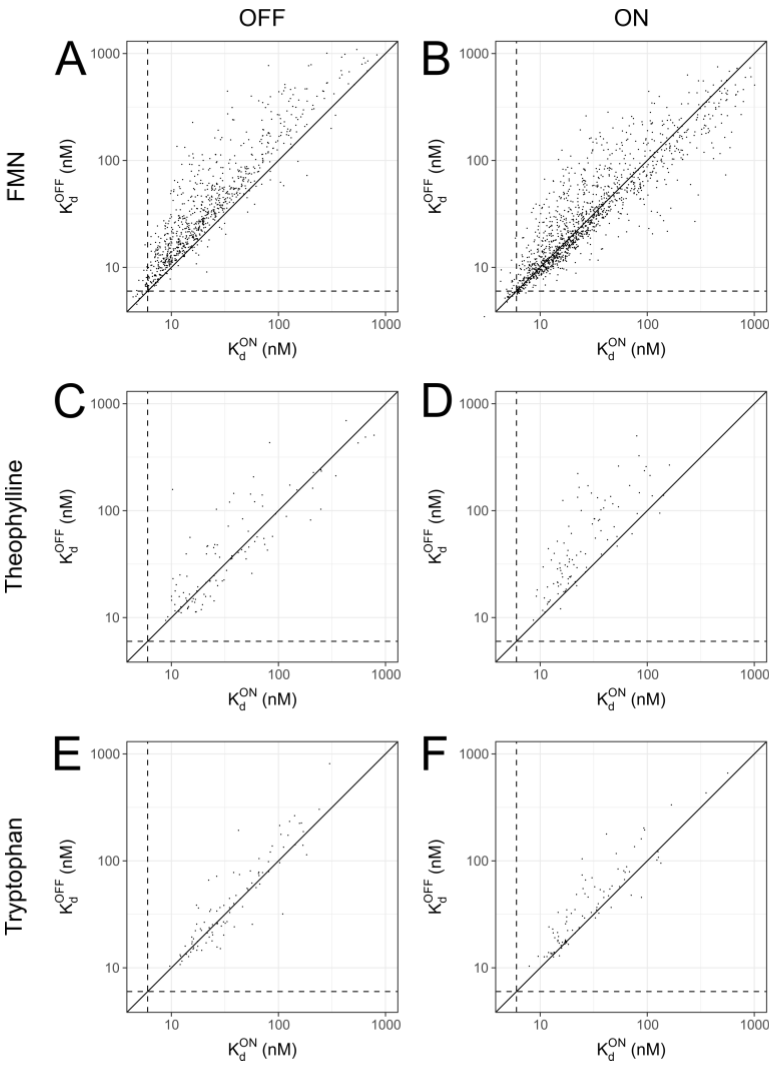

### Figure S3

**Number of iterations to convergence.** The number of iterations of Monte Carlo to reach constraint satisfaction varied across different ligands. On average, every 1,000 iterations took about 2 minutes on one core.

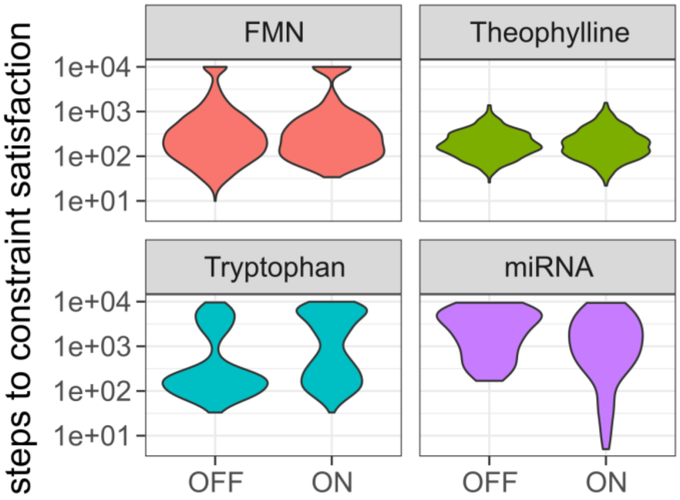

### Figure S4

**Secondary structure features and activation ratio.** The data show that some secondary structure features correlate significantly (Fisher z-transform) with activation ratio. These include the number of bulges and number of hairpin/internal/multi loops in the absence of ligand as well as the number of internal loops in the presence of ligand. Further, more shared base pairs between states was correlated with higher activation ratios.

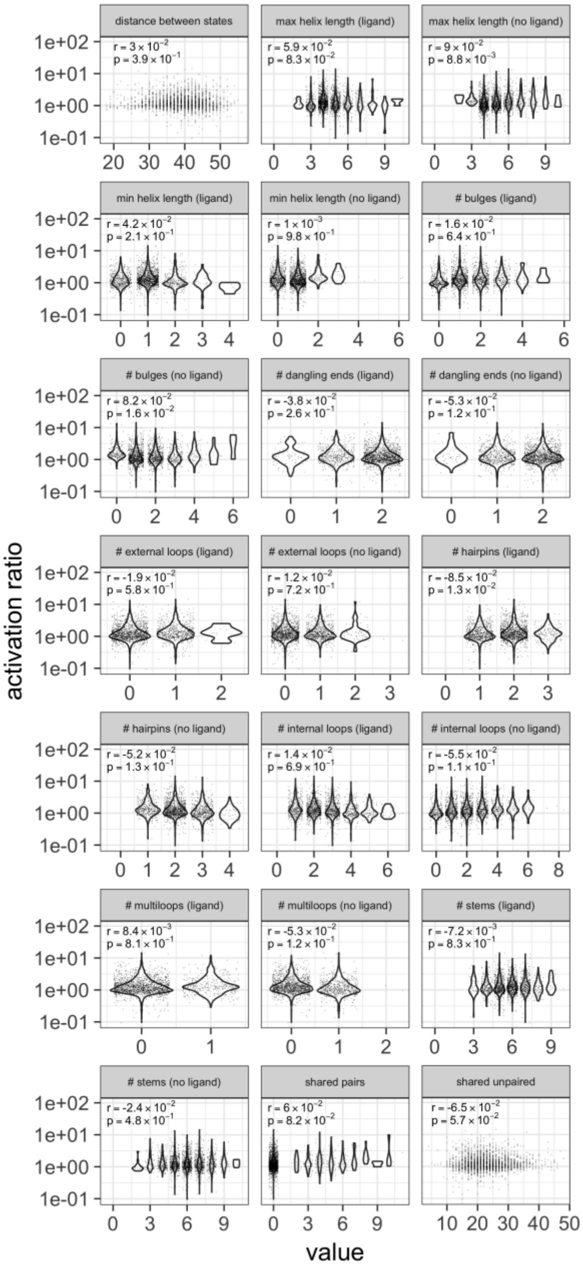

### Figure S5

**Comparison of predicted and measured $K_d$ values.** For both small molecule (A) and miRNA (B) riboswitches, there is a significant correlation between predicted and measured $K_d$ values, but the degree of correlation is poor.

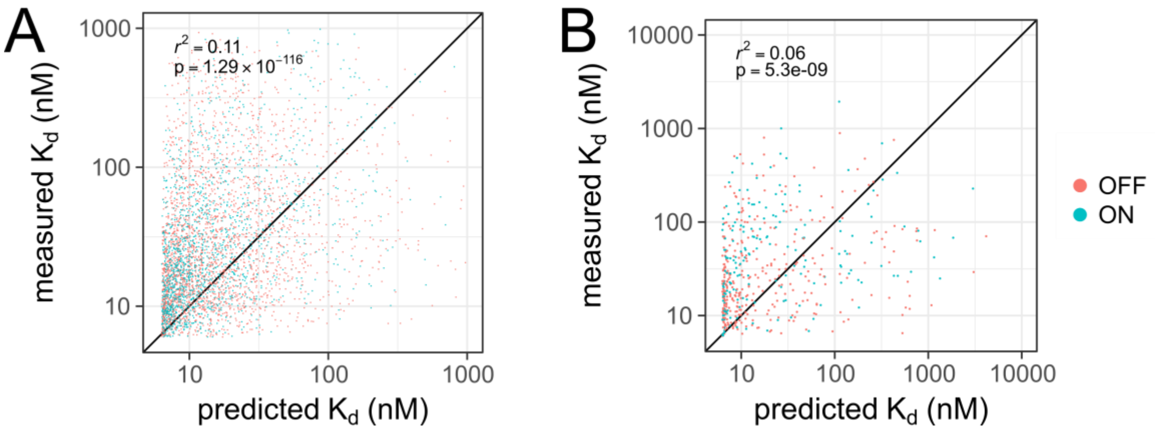

### Table S1

**Summary statistics for activation ratios for RiboLogic and baseline.**

| Design | RiboLogic | | | | baseline | | | |
|---|---|---|---|---|---|---|---|---|
| | max | median | standard deviation | count | max | median | standard deviation | count |
| FMN OFF | 9.74 | 0.987 | 0.878 | 1357 | 1.15 | 0.869 | 0.144 | 524 |
| FMN ON | 14.4 | 1.46 | 1.33 | 849 | 2.90 | 1.15 | 0.308 | 524 |
| theophylline OFF | 9.92 | 1.73 | 1.60 | 97 | 4.62 | 1.22 | 0.369 | 366 |
| theophylline ON | 15.4 | 0.991 | 1.65 | 99 | 1.33 | 0.820 | 0.155 | 366 |
| tryptophan OFF | 4.29 | 1.17 | 0.632 | 89 | 2.47 | 1.01 | 0.206 | 392 |
| tryptophan ON | 4.55 | 1.08 | 0.576 | 94 | 1.97 | 0.988 | 0.181 | 392 |
| miRNA OFF | 21.8 | 0.825 | 1.68 | 188 | 4.95 | 0.819 | 0.483 | 188 |
| miRNA ON | 20.0 | 1.17 | 2.20 | 98 | 4.26 | 1.23 | 0.584 | 98 |

### Table S2

**Summary of statistical tests comparing RiboLogic activation ratios.** All comparisons used a two-sided Wilcoxon rank sum test.

| design | RiboLogic vs baseline | RiboLogic vs non-switching (activation ratio 1) |
|---|---|---|
| FMN OFF | $8.0 \times 10^{-41}$ | $1.1 \times 10^{-6}$ |
| FMN ON | $8.5 \times 10^{-58}$ | $2.8 \times 10^{-130}$ |
| Theophylline OFF | $3.5 \times 10^{-13}$ | $3.0 \times 10^{-16}$ |
| Theophylline ON | $2.0 \times 10^{-10}$ | 0.071 |
| Tryptophan OFF | $3.4 \times 10^{-13}$ | $1.8 \times 10^{-10}$ |
| Tryptophan ON | $6.0 \times 10^{-14}$ | $1.5 \times 10^{-3}$ |
| miRNA OFF | 0.98 | $7.4 \times 10^{-7}$ |
| miRNA ON | 0.15 | $1.7 \times 10^{-4}$ |

### Table S3

**Summary of best-of-ten analysis.** All values are based on 1,000 bootstrap samples of 10 designs each. Comparisons were made using a two-sided Wilcoxon rank sum test.

| design | RiboLogic median | Baseline median | p-value |
|---|---|---|---|
| FMN OFF | 2.57 | 1.01 | $< 2 \times 10^{-308}$ |
| FMN ON | 3.89 | 1.69 | $1.8 \times 10^{-252}$ |
| Theophylline OFF | 4.86 | 1.72 | $3.3 \times 10^{-281}$ |
| Theophylline ON | 3.44 | 1.04 | $< 2 \times 10^{-308}$ |
| Tryptophan OFF | 2.28 | 1.24 | $2.4 \times 10^{-264}$ |
| Tryptophan ON | 2.08 | 1.23 | $3.7 \times 10^{-237}$ |
| miRNA OFF | 1.66 | 1.52 | $8.5 \times 10^{-6}$ |
| miRNA ON | 2.84 | 2.10 | $7.2 \times 10^{-26}$ |
